# Convergent Met and voltage-gated Ca^2+^ channel signaling on Ras-Erk MAPK drives migratory activation of dendritic cells parasitized by *Toxoplasma gondii*

**DOI:** 10.1101/2020.01.08.898197

**Authors:** Einar B. Ólafsson, Arne L. ten Hoeve, Xiaoze Li Wang, Linda Westermark, Manuel Varas-Godoy, Antonio Barragan

**Author notes:** Correspondence to Antonio Barragan, Mailing address: Department of Molecular Biosciences, The Wenner-Gren Institute, Stockholm University, 106 91 Stockholm, Sweden.

## Abstract

Ras-Erk MAPK signaling controls many of the principal pathways involved in metazoan cell motility, drives metastasis of multiple cancer types and is targeted in chemotherapy. Yet, its putative roles in immune cell functions or in infections have remained elusive. Here, using primary dendritic cells (DCs) in an infection model with the protozoan *Toxoplasma gondii*, we show that two pathways activated by infection converge on Ras-Erk MAPK signaling to promote migration of parasitized DCs. We identify signaling through the receptor tyrosine kinase Met (also known as HGFR) as a driver of *T. gondii*-induced DC hypermotility. Further, we show that voltage-gated Ca^2+^channel (VGCC, subtype Ca_V_1.3) signaling impacts the migratory activation of DCs via calmodulin-calmodulin kinase II. We report that VGCC and Met signaling converge on Ras GTPase to drive Erk1/2 phosphorylation and migratory activation of *T. gondii*-infected DCs. The data provide a molecular basis for the hypermigratory mesenchymal-to-amoeboid transition (MAT) of parasitized DCs. The emerging concept suggests that parasitized DCs acquire metastasis-like migratory properties to promote infection-related dissemination.

## Introduction

The mitogen-activated protein kinase (MAPK) Ras-Erk signaling axis plays important roles in cell migration, cancer metastasis, cell proliferation and survival (Eblen, 2018). Ras GTPases function as molecular switches by cycling between an inactive and active state. Their function in cell motility is linked to localization to the plasma membrane, which is mediated by farnesylation (Cox et al., 1992). At the plasma membrane, Ras is locally activated by guanine-nucleotide exchange factors (GEFs). In turn, Ras activates the Raf-Mek-Erk signaling cascade which regulates cell migration (Chernyavsky et al., 2005; Shi et al., 2019). In proximity of the cell membrane, Ras activation can be mediated by diverse mechanisms, including receptor tyrosine kinase (RTK) signaling, integrin activation and calcium (Ca^2+^) influx (Giehl et al., 2000; Rosen et al., 1994; Schlaepfer et al., 1994).

In cancer cells, the RTK Met (also termed c-Met or hepatocyte growth factor receptor/HGFR) promotes cell migration and metastasis upon binding its ligand, hepatocyte growth factor (Hgf, Scatter Factor/SF)(Webb et al., 1998). Activated Met recruits Ras-GEFs, which locally activate Ras, leading to Erk phosphorylation and cell motility (Hartmann et al., 1994; Wang et al., 2002). Additionally, voltage-gated Ca^2+^ channels (VGCCs) mediate Ca^2+^ influx which activates Ca^2+^-sensing proteins locally at the plasma membrane (Dolmetsch et al., 2001; Rosen et al., 1994). In the context of cell migration, Ca^2+^ influx activates calmodulin (CaM) and downstream kinases, which impacts cytoskeleton organization and cell motility by regulating actin dynamics through the Ras-Erk pathway (Li et al., 2004; Lundberg et al., 1998). Despite the central roles played by MAPK signaling in immune responses (Stupack et al., 2000), the impact of Ras-Erk signaling on the diverse immune cell functions has only started to be understood (Ebert et al., 2016; Scheele et al., 2007). Further, in primary dendritic cells (DCs), Ras-Erk signaling remains largely unexplored (Riegel et al., 2019).

Owing to host-pathogen coevolution with reciprocal selection, the study of host-pathogen interactions has emerged as a powerful approach to gain insight into basic cell biology. The protozoan *Toxoplasma gondii* is a model obligate intracellular pathogen due to its wide host range among warm-blooded vertebrates and ability to actively invade nucleated cells (Sibley, 2004).

One third of the global human population is estimated to be chronically infected by *T. gondii* (Pappas et al., 2009). Upon ingestion and after crossing the intestinal epithelium, *T. gondii* tachyzoites rapidly disseminate, ultimately establishing chronic infection in the brain (Montoya and Liesenfeld, 2004). Early on, tachyzoites encounter DCs, which play a determinant role in mounting a robust protective immune response (Liu et al., 2006). Paradoxically, *T. gondii* exploits the inherent migratory ability of DCs for dissemination via a ‘Trojan horse’ mechanism (Courret et al., 2006; Lambert et al., 2006; Lambert et al., 2009; Sangare et al., 2019). Within minutes of active invasion by *T. gondii*, DCs adopt a hypermigratory phenotype that mediates rapid systemic dissemination in mice (Kanatani et al., 2017; Weidner and Barragan, 2014). We recently reported that *T. gondii* induces mesenchymal-to-amoeboid transition (MAT) in DCs (Olafsson et al., 2018). Hypermigratory parasitized DCs are characterized by high-velocity amoeboid locomotion (termed hypermotility), cytoskeletal reorganization with loss of podosomes, decreased pericellular proteolysis and redistribution of integrins (Kanatani et al., 2015; Olafsson et al., 2018; Weidner et al., 2013). Hypermigration is triggered by non-canonical GABA_A_ receptor-mediated (Fuks et al., 2012) activation of VGCCs, subtype Ca_V_1.3 (Kanatani et al., 2017) and Timp-1-mediated Itgb1-dependent activation of focal adhesion kinase (Fak) (Olafsson et al., 2019). However, the signaling elements downstream of VGCC and Itgb1-Fak signaling, and their respective contribution to the hypermigratory phenotype of DCs, have remained unknown.

Here, we use primary DCs and *T. gondii* to investigate alternative pathways for migratory activation of immune cells. We identify Ras-dependent sustained Erk phosphorylation downstream of VGCC-CaM-CaMkII signaling and Met signaling, with an impact on non-canonical migratory activation of *T. gondii*-infected DCs. The findings provide novel mechanistic insights into infection-related MAT of DCs.

## Results

### Erk is rapidly phosphorylated in parasitized DCs and gene silencing of *erk1* and *erk2* abolishes DC hypermotility

Because the MAPK Erk1/2 (Erk) plays a key role in cancer cell metastasis (Viala and Pouyssegur, 2004) and leukocyte motility (Stupack et al., 2000), we investigated Erk activity in primary murine DCs challenged with *T. gondii* tachyzoites. First, we found that infection impacted *erk1* and *erk2* mRNA expression (Fig. S1A). However, non-significant differences in total Erk protein expression were detected upon *T. gondii* challenge 6 hours post-infection (hpi) (Fig. S1B). In sharp contrast, elevated Erk phosphorylation was observed at this timepoint, which was abrogated upon treatment with the Mek inhibitor u0126 (Fig. 1A). Because DC hypermigration sets in within minutes of *T. gondii* invasion (Weidner et al., 2013), we assessed the early kinetics of Erk phosphorylation upon *T. gondii* challenge. As we found that the ratio of phosphorylated Erk to total Erk was rapidly elevated in *T. gondii*-challenged DCs (Fig. 1B), we investigated its putative role in DC hypermotility. Importantly, inhibition of Erk phosphorylation by Mek antagonism significantly decreased the mean velocities of *T. gondii*-infected DCs, with a non-significant impact on the base-line motility of DCs (Fig. 1C; Fig. S1C). Mek antagonism non-significantly impacted DC invasion by *T. gondii* (Fig. S1D, E).

**Figure 1.**
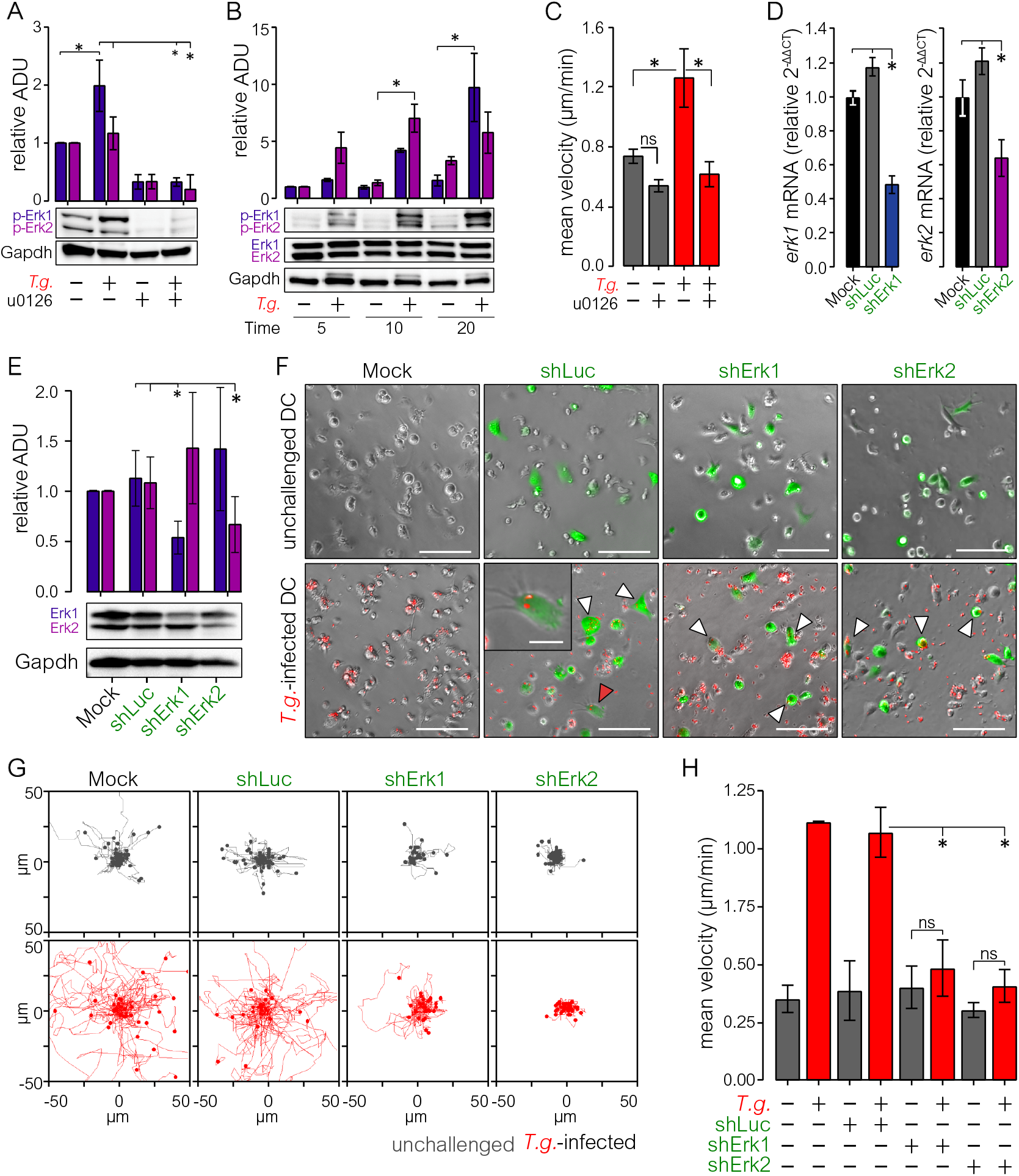
Rapid phosphorylation of Erk upon *T. gondii* infection and its impact on DC hypermotility. **A.** Erk phosphorylation in unchallenged and *T. gondii*-challenged DCs (*T.g*.) ± u0126 treatment at 6 h post-infection (hpi). Bar graph shows mean (± SEM) total protein related to Gapdh (relative arbitrary density units, ADU). Expression of unchallenged DCs was set to 1. Images show representative blots used for quantification from 1 experiment (n=8 biological replicates from independent blots). **B.** Erk phosphorylation in unchallenged and *T. gondii*-challenged DCs (*T.g.*) at indicated time points. Bar graph shows mean (± SEM) phosphorylated protein related to Gapdh. Expression of unchallenged DCs was set to 1. Images show representative blots of total and phosphorylated Erk and Gapdh used for quantification (n=4 biological replicates from independent blots). **C.**Bar graph shows mean (± SEM) velocity of unchallenged and *T. gondii*-infected DCs (*T.g.*) embedded in collagen ± u0126 treatment (10 μM; n=3 independent experiments). **D.** Relative erk1 and erk2 mRNA expression (relative 2-ΔCt) of mock-treated, control shLuc- and shErk1- and shErk2-transduced DCs. Expression of mock-treated DCs was set to 1 (n=3 independent experiments). **E.** Relative Erk1 and Erk2 total protein expression of DCs transduced with control shLuc or shErk1 or shErk2 lentivirus. Expression of mock-treated DCs was set to 1. Bar graph shows mean (± SEM) Erk1 and Erk2 protein expression related to Gapdh. Blots show representative blots used for quantification (n=3 biological replicates from independent blots). **F.** Representative micrographs of mock-treated DCs (Mock) and eGFP-expressing DCs transduced with lentiviral vectors targeting erk1 and erk2 mRNA (shErk1 and shErk2, respectively) or a non-expressed target (shLuc) embedded in collagen ± RFP-expressing *T. gondii* tachyzoites (*T.g.*). Arrowheads indicate *T. gondii*-infected cells expressing eGFP assessed in the assay. Scale bars: 200 μm. Inset image shows representative transduced DC (green) infected by *T. gondii* (red), indicated by a red arrowhead. Scale bar: 50 μm. Micrographs are representative of 3 independent experiments. **G.** Representative motility plots of unchallenged and *T. gondii*-infected (*T.g.*) mock-treated, shLuc-, shErk1- and shErk2-transduced DCs embedded in collagen. Plots are representative of 3 independent experiments. **H.** Mean velocity of unchallenged and *T. gondii*-infected (*T.g.*) mock-treated, shLuc-, shErk1- and shErk2-transduced DCs embedded in collagen (n=3 independent experiments). Asterisks (*) indicate significant difference, ns: non-significant difference: One-way ANOVA, Holm-Sidak’s multiple comparisons test (A, C-D and H). Two-way ANOVA, Holm-Sidak’s multiple comparisons test (B). Paired *t*-test (E).

To determine the respective roles of Erk1 and Erk2 in DC hypermotility, we transduced DCs with lentivirus encoding an eGFP reporter and shRNA targeting *erk1* (shErk1) or *erk2* (shErk2) mRNA or non-expressed mRNA (shLuc). First, the knock down efficiency of shErk1 and shErk2 was quantified. ShErk1- and shErk2-transduced DCs exhibited significantly reduced levels of Erk1 and Erk2 mRNA (Fig. 1D) and protein (Fig. 1E), respectively, related to mock/shLuc-transduced DCs. Next, the motility of transduced parasitized DCs was analyzed (Fig. 1F, G). Importantly, gene silencing of both *erk1* and *erk2* significantly reduced hypermotility in *T. gondii*-infected DCs and had a non-significant impact on baseline motility (Fig. 1H). From these data, we conclude that challenge of DCs with *T. gondii* is accompanied by a rapid phosphorylation of Erk that underlies *T. gondii*-mediated migratory activation of DCs.

### Erk phosphorylation via Ras downstream of VGCC-CaM-CaMkII-mediated Ca^2+^ signaling governs DC hypermotility

Based on concepts in neuronal cells showing that Erk is activated downstream of VGCC signaling (Kotturi et al., 2003) and because VGCCs are implicated in *T. gondii*-mediated hypermotility (Kanatani et al., 2017), we hypothesized that VGCC-Ras-Raf-Mek signaling mediated Erk phosphorylation in *T. gondii*-infected DCs. To this end, we analyzed the phosphorylation of Erk in *T. gondii*-infected DCs upon treatment with the VGCC inhibitors nifedipine (targeting L-type VGCCs) and CPCPT (a selective inhibitor of VGCC subtype Ca_V_1.3). Interestingly, a significant inhibition of Erk phosphorylation was observed upon VGCC antagonism with both nifedipine and CPCPT in *T. gondii*-infected DCs (Fig. 2A). VGCC inhibition non-significantly impacted on baseline Erk phosphorylation in unchallenged DCs (Fig. S2A). Because motility-related Ras signaling is strongly associated to localized farnesylated-Ras activity at the plasma membrane (Charest et al., 2010; Sasaki et al., 2004), we pharmacologically blocked plasma membrane Ras activity. Hence, the Ras farnesylation inhibitor salirasib significantly inhibited Erk phosphorylation in *T. gondii*-infected DCs (Fig. 2B), with a non-significant impact on baseline Erk phosphorylation (Fig. S2B) or *T. gondii* invasion (Fig. S2C). Conversely, VGCC agonism with Bay K, a structural analog of nifedipine with a positive ionotropic effect, led to elevated Erk phosphorylation, which was inhibited by salirasib treatment (Fig. 2C). Importantly, hypermotility was abrogated by Ras antagonism (Fig. 2D, E), in line with VGCC inhibition, Ca_V_1.3 gene silencing (Kanatani et al., 2017) and Mek inhibition (Fig. 1C; Fig. S1D), all indicating the implication of a VGCC-Ras-Mek signaling axis.

**Figure 2.**
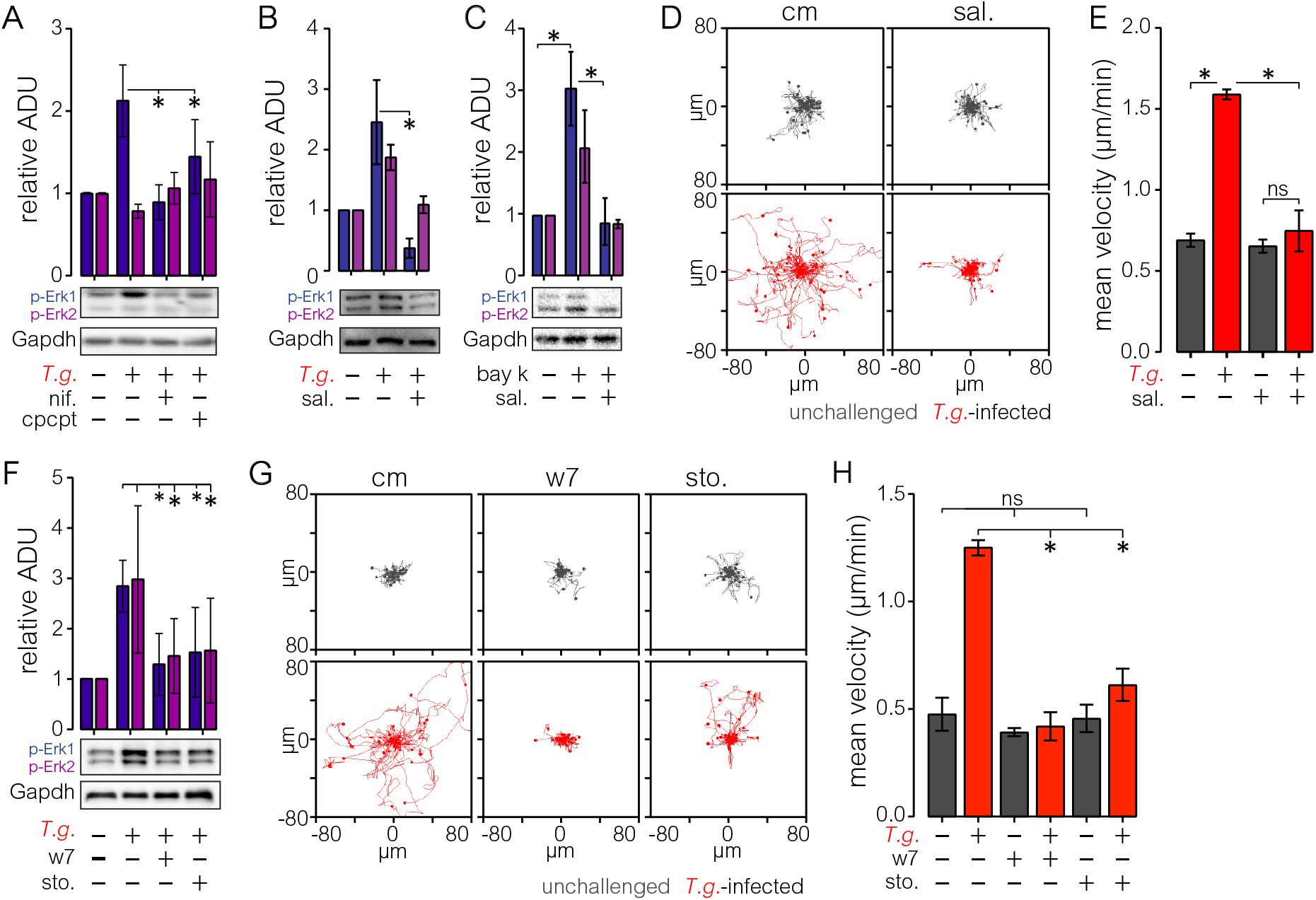
Activation of VGCCs mediates Erk phosphorylation through CaM-CaMkII-Ras and impacts DC motility. **A.** Erk phosphorylation in unchallenged and *T. gondii*-challenged DCs (*T.g.*) ± nifedipine (nif., 50μM) or CPCPT (10 μM) treatment 2 hpi. Bar graph shows mean (± SEM) phosphorylated Erk1 and Erk2 protein related to Gapdh. Expression of unchallenged DCs was set to 1. Images show representative blots used for quantification (n=3 biological replicates from independent blots). **B.** Erk phosphorylation in unchallenged and *T. gondii*-challenged DCs (*T.g.*) ± salirasib (sal., 100 μM) 2 hpi. Bar graph shows mean (± SEM) phosphorylated Erk1 and Erk2 protein related to Gapdh. Expression of unchallenged DCs was set to 1. Images show representative blots (n=3 biological replicates from independent blots). **C.** Erk phosphorylation in unchallenged DCs ± Bay K (10 μM) and salirasib (sal., 100 μM) 20 min post-infection. Bar graph shows mean (± SEM) phosphorylated Erk1 and Erk2 protein related to Gapdh. Expression of unchallenged DCs was set to 1. Representative blots are shown (n=3 biological replicates from independent blots). **D.** Representative motility plots of unchallenged and *T. gondii-infected* DCs (*T.g.*) embedded in collagen ± salirasib (sal., 25 μM). Plots are representative of 3 independent experiments. **E.** Bar graph shows mean velocity (± SEM) of unchallenged and *T. gondii-infected* DCs (*T.g.*) embedded in collagen treated as in D (n=3 independent experiments). **F.** Erk phosphorylation in unchallenged and *T. gondii*-challenged DCs (*T.g.*) ± w7 (25 μM) or sto-609 (sto., 50 μM) treatment 2 hpi. Bar graphs show mean (± SEM) phosphorylated Erk1 and Erk2 protein related to Gapdh. Expression of unchallenged DCs was set to 1. Images show representative blots (n=4 biological replicates from independent blots). **G.** Representative motility plots of unchallenged and *T. gondii*-infected DCs (*T.g.*) embedded in collagen ± w7 (25 μM) or sto-609 (sto., 25 μM). Plots are representative of 3 independent experiments. **H.** Bar graph shows mean velocity (± SEM) of unchallenged and *T. gondii*-infected DCs (*T.g.*) embedded in collagen treated as in G (n=3 independent experiments). Asterisks (*) indicate significant difference, ns: non-significant difference: One-way ANOVA, Tukey’s HSD post-hoc test (A-C, E and H), Holm-Sidak’s multiple comparisons test (F)

Next, because calmodulin (CaM) and CaM kinase II (CaMkII) act downstream of VGCC-mediated Ca^2+^ influx and impact Ras-Raf activity and Erk phosphorylation (Agell et al., 2002), we investigated the putative roles of CaM and CaMkII in *T. gondii*-mediated Erk phosphorylation and DC motility. Clearly, antagonism of CaM and CaMkII significantly reduced *T. gondii*-induced Erk phosphorylation (Fig. 2F). However, a significant reduction in Erk phosphorylation was also observed in unchallenged DCs upon CaM inhibition, but not CaMkII inhibition (Fig S2B), indicating implication of CaM in baseline phosphorylation of Erk. Non-significant effects were observed on *T. gondii* invasion (Fig. S2C). Importantly, CaM and CaMkII antagonism inhibited hypermotility of *Toxoplasma-infected* DCs (Fig. 2G, H). A non-significant reduction of baseline motility was also noted in unchallenged DCs upon CaM inhibition, but not CaMkII inhibition, (Fig. 2G, H), likely consistent with a broader impact of CaM on intracellular Ca^2+^ homeostasis and a more selective effect of CaMkII on Ras and Raf (Fig S2B) (Illario et al., 2003).

Together, these data indicate that L-type VGCC-Ca_V_1.3 signal transduction occurs via the CaM-CaMkII-Ras pathway in parasitized DCs, with subsequent phosphorylation of Erk and an impact on DC migration.

### Hgf-Met signaling promotes DC motility via Ras activation and Erk phosphorylation

We recently reported the implication of the non-receptor tyrosine kinase Fak in DC hypermotility (Olafsson et al., 2019). However, receptor tyrosine kinases (RTKs) also play central roles in cell migration and metastasis (Lemmon and Schlessinger, 2010). Because aberrantly activated RTK Met mediates metastasis in a number of cancers (Benvenuti and Comoglio, 2007; Webb et al., 1998), we investigated the expression of Met and its ligand Hgf in DCs challenged with *T. gondii*. Kinetics analyses showed that DCs upregulated the transcription of *met* and *hgf* mRNAs shortly after *T. gondii* challenge (Fig. S3A, B), motivating further functional investigation.

First, we investigated the impact of Met activation with recombinant Hgf (rHgf) on Erk phosphorylation and DC motility. Indeed, treatment with rHgf induced Erk phosphorylation in unchallenged DCs (Fig. 3A). Further, rHgf treatment stimulated motility in both unchallenged and *T. gondii*-challenged DCs (Fig. 3B). Importantly, *T. gondii*-induced Erk phosphorylation was significantly inhibited upon treatment with the Met inhibitor su11274 (targeting submembrane Met phosphorylation) and was not rescued by rHgf treatment (Fig. 3C). Moreover, inhibition of Met phosphorylation inhibited DC hypermotility, with a non-significant impact on DC baseline motility (Fig. 3D, E). Unexpectedly, challenge with *T. gondii* non-significantly impacted the amounts of secreted Hgf in supernatants, related to unchallenged DCs (Fig. 3F). Because Ras acts downstream of Met in its activation of Erk (Wang et al., 2002) and Ras inhibition inhibited hypermotility (Fig. 2E) and Erk phosphorylation (Fig. 2B), we investigated the impact of salirasib treatment on rHgf-mediated Erk phosphorylation. Similar to its inhibitory effect on VGCC-mediated Erk phosphorylation, salirasib treatment significantly inhibited rHgf-mediated Erk phosphorylation (Fig. 3G).

**Figure 3.**
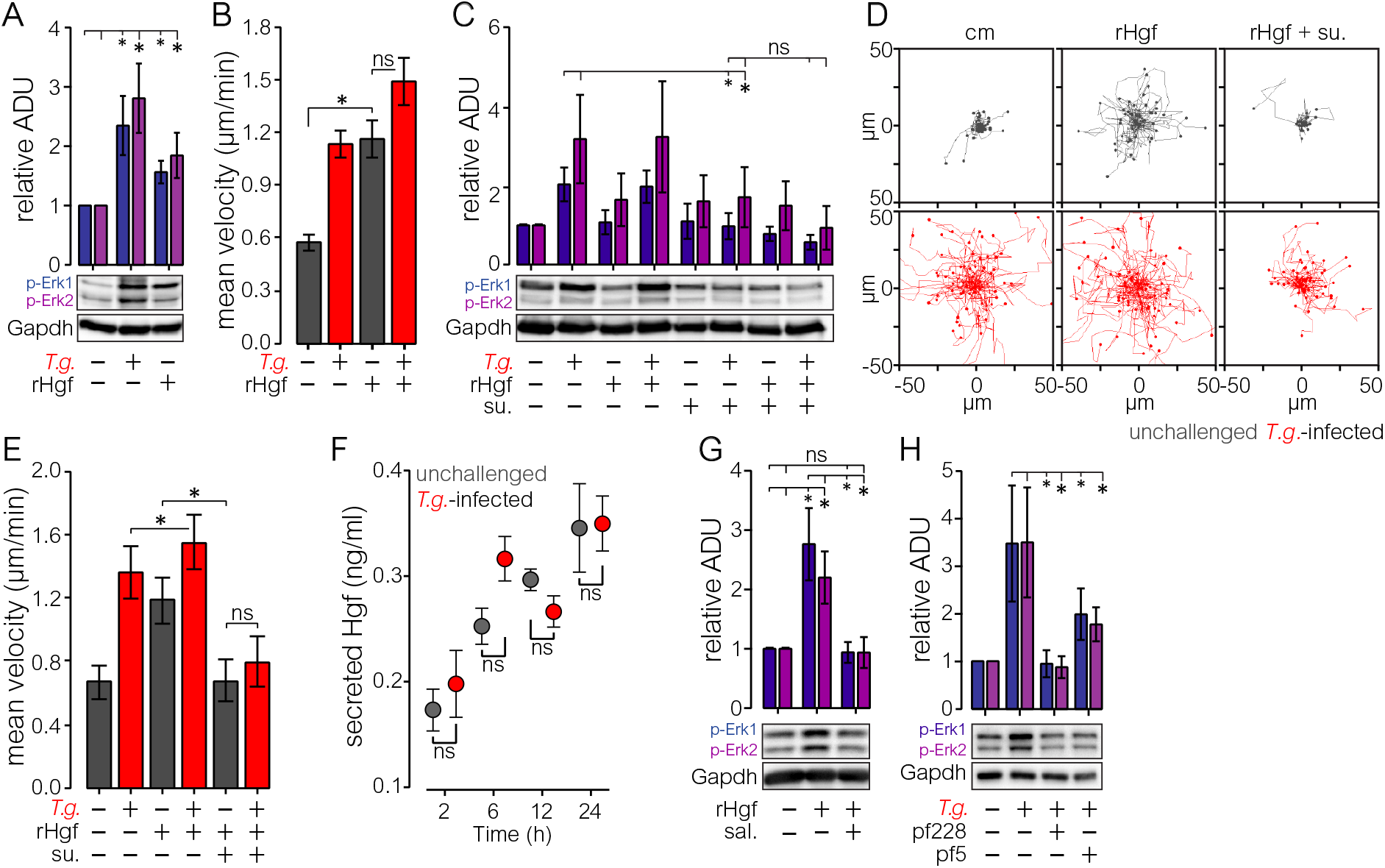
Erk phosphorylation downstream of Hgf-Met via Ras signaling impacts DC hypermotility. **A.** Erk phosphorylation in unchallenged and *T. gondii*-challenged DCs (*T.g.*) ± recombinant-Hgf (rHgf, 40 ng/ml) treatment 2 hpi. Bar graph shows mean (± SEM) phosphorylated protein related to Gapdh. Expression of unchallenged DCs was set to 1. Images show representative blots used for quantification from 1 experiment (n=6 biological replicates from independent blots). **B.** Mean velocity of unchallenged and *T. gondii*-infected DCs (*T.g.*) embedded in collagen ± rHgf (40 ng/ml) treatment. Plots are representative of 3 independent experiments. **C.** Erk phosphorylation in unchallenged and *T. gondii-challenged* DCs (*T.g.*) ± rHgf (40 ng/ml), su11274 (10 μM) or rHgf + su11274 treatment 6 hpi. Bar graph shows mean (± SEM) phosphorylated protein related to Gapdh. Expression of unchallenged DCs was set to 1. Images show representative blots (n=4 biological replicates from independent blots). **D.**Representative motility plots of unchallenged and *T. gondii*-infected DCs (*T.g.*) embedded in collagen ± rHgf (40 ng/ml), su11274 (10 μM) or rHgf + su11274 treatment. Plots are representative of 3 independent experiments. **E.** Bar graph shows mean velocity (± SEM) of unchallenged and *T. gondii*-infected DCs (*T.g.*) embedded in collagen treated as in D (n=3 independent experiments). **F.** Secreted Hgf protein in supernatants from unchallenged and *T. gondii*-challenged DCs (*T.g.*) collected 2, 6, 12 and 24 hpi. Data are presented as mean (± SEM) from 3 experiments (n=3). **G.** Erk phosphorylation in unchallenged DCs ± rHgf (40 ng/ml) or rHgf and salirasib (sal., 100 μM) 20 min. post infection. Bar graph shows mean (± SEM) phosphorylated Erk1 and Erk2 protein related to Gapdh. Expression of unchallenged DCs was set to 1. Images show representative blots (n=4 biological replicates from independent blots). **H.** Erk phosphorylation in unchallenged and *T. gondii*-challenged DCs (*T.g.*) ± pf228 (10 μM) or pf5 (10 μM) treatment 2 hpi. Bar graph shows mean (± SEM) phosphorylated protein related to Gapdh. Expression of unchallenged DCs was set to 1. Images show representative blots (n=3 biological replicates from independent blots). Asterisks (*) indicate significant difference, ns: non-significant difference: One-way ANOVA, Tukey’s HSD post-hoc test (A, B, E and G-H). Paired t-test (C). Two-way ANOVA, Holm-Sidak’s multiple comparisons test (F).

In absence of elevated Hgf secretion upon *T. gondii* challenge and because *T. gondii* activates integrin Itgb1-Fak signaling in DCs (Olafsson et al., 2019), we hypothesized that, in addition to Hgf, Hgf-independent activation of Met might occur via Fak (Hui et al., 2009). Indeed, Erk phosphorylation was significantly reduced in *T. gondii*-infected DCs upon treatment with inhibitors targeting Fak and the associated kinase Pyk2 (Fig. 3H). The treatments impacted non-significantly on Erk phosphorylation in unchallenged DCs (Fig. S3C), on parasite invasion and on infection frequency (Fig. S3D).

Taken together, the data show that Met-Ras signaling is activated in DCs upon *T. gondii* infection, with an impact on motility via phosphorylation of Erk. Further, our data suggest that, in parasitized DCs, Met is activated by both its ligand Hgf and via tyrosine kinase-mediated transactivation.

### Gene silencing of *met* inhibits hypermotility of *T. gondii-infected* DCs

To determine the role of Met in *T. gondii*-induced hypermigration, we applied a gene silencing approach in primary DCs. First, we analyzed Met protein expression in DCs by flow cytometry (Fig. S4A, B). Consistent with *met* transcription data, signal corresponding to Met protein was significantly elevated in CD11c^+^ DCs infected by *T. gondii* (Fig. 4A). Of note, parasitized DCs exhibited significantly higher signal corresponding to plasma membrane-associated Met and total Met, compared with non-infected by-stander DCs (Fig. 4A). This indicated a selective upregulation of Met in parasite-invaded DCs.

**Figure 4.**
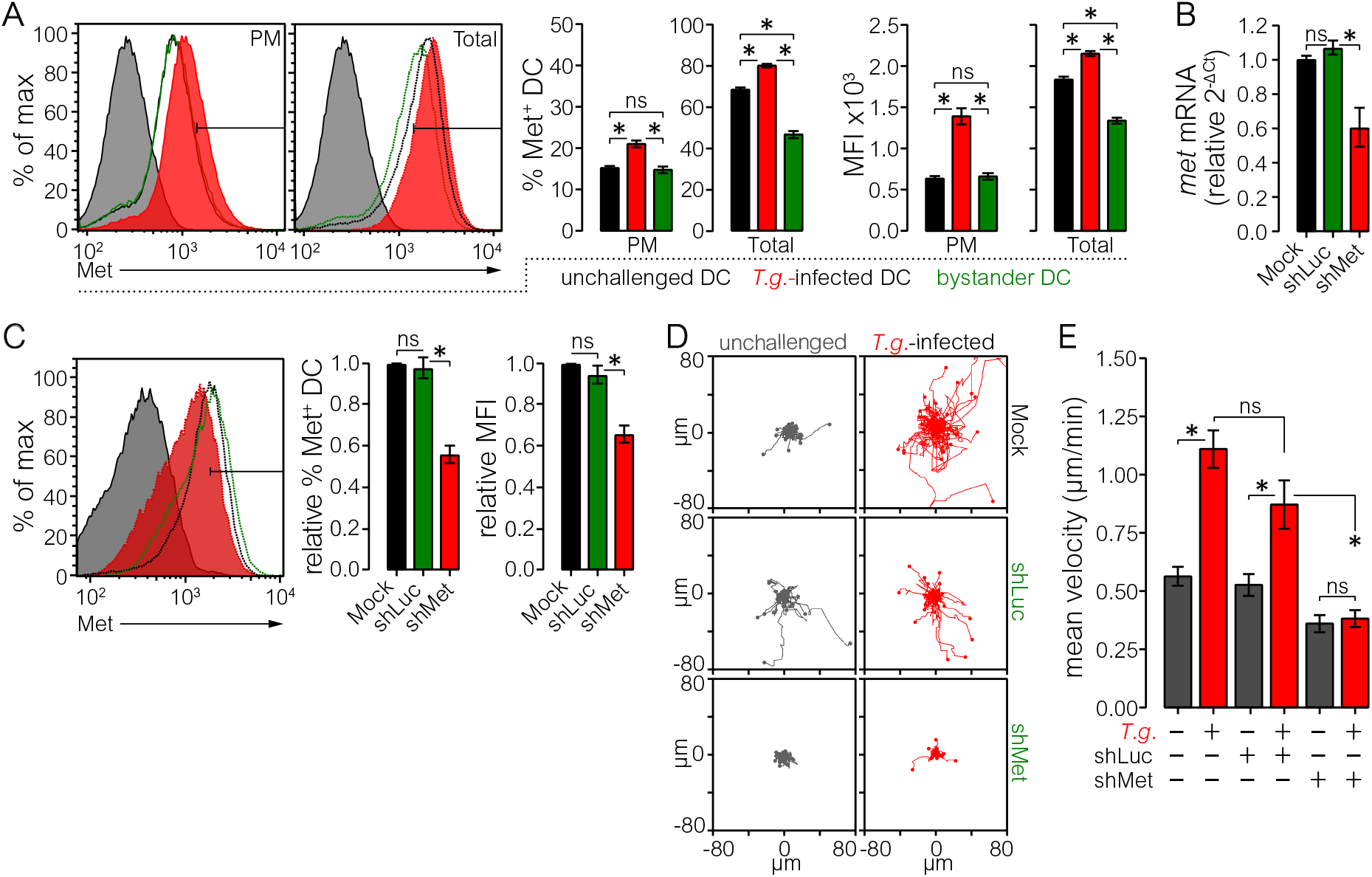
Expression of Met is upregulated in parasitized DCs and gene silencing of met inhibits hypermotility. **A.** Plasma membrane (PM, left histogram) and total (right histogram) Met protein expression of CD11c^+^ unchallenged (black), non-infected bystander (green) and *T. gondii*-infected (*T.g.*, red) DCs. Histograms show gate used to define Met^+^ cells. Dashed and continuous lines show total and PM Met expression, respectively. Grey fill represents unstained control for CD11c and isotype control for Met. Bar graphs show the percentage of Met^+^ cells and Met median fluorescence intensity (MFI) for each population (n=4 independent experiments). **B.** Relative Met mRNA expression (2^-ΔCt^) of mock-treated, control shLuc- and shMet-transduced DCs. Expression of mock-treated DCs was set to 1 (n=3 independent experiments). **C.** Met protein expression of CD11c^+^ mock-treated DCs and DCs transduced with control shLuc or shMet lentivirus. Gating performed as in A. Histogram shows gate used to define Met^+^ cells. Black, Green and Red show mock-treated, shLuc-transduced and shMet-transduced DCs, respectively. Grey fill represents isotype control. Bar graphs show the percentage of Met^+^cells and Met MFI for each population (n=3 independent experiments). **D.** Representative motility plots of unchallenged and *T. gondii*-infected (*T.g.*) mock-treated, shLuc-, shMet-transduced DCs embedded in collagen. Plots are representative of 3 independent experiments. **E.** Mean velocity (± SEM) of unchallenged and *T. gondii*-infected (*T.g.*) mock-treated, shLuc- or shMet-transduced DCs embedded in collagen (n=6 independent experiments). Asterisks (*) indicate significant difference, ns: non-significant difference: One-way ANOVA, Tukey’s HSD post-hoc test (A-C and E).

Next, we silenced *met* mRNA expression with lentivirus encoding an eGFP reporter and shRNA targeting *met* mRNA (shMet) or non-target control (shLuc). First, we confirmed that DCs transduced with shMet exhibited significantly reduced *met* mRNA (Fig. 4B, Fig S4C) and Met total protein (Fig. 4C), relative to shLuc transduced DCs. Finally, Met-silenced *T. gondii*-challenged DCs were subjected to motility analyses (Fig S4D). Importantly, gene silencing of *met* abolished hypermotility in *T. gondii*-infected DCs (Fig. 4D, E), in line with data of pharmacological antagonism of Met (Fig 3E). Altogether, these data show that Met signaling is activated in *T. gondii*-infected DCs with an impact on motility.

## Discussion

In the present study we investigated the putative role of MAPK Erk1/2 (Erk) in *T. gondii*-induced hypermigration of primary DCs. We report that *T. gondii*-induced VGCC and Met signaling converge on the Ras-Erk signaling axis to mediate hypermotility of parasitized DCs.

Our studies establish that Erk signaling plays a central role for *Toxoplasma*-induced hypermigration of DCs. A rapid and sustained Erk phosphorylation was observed upon *T. gondii* challenge, leading to the activation of DC motility. Despite a transcriptional impact by the infection, total Erk protein amounts remained stable, likely due to the 50 h half-life time of Erk (Schwanhausser et al., 2011). By shRNA-mediated gene silencing of *erk1* and *erk2* and pharmacological antagonism of Erk phosphorylation (by Mek inhibition), we demonstrate that phosphorylated Erk1 and Erk2 are necessary for DC hypermotility. Interestingly, knock down of Erk1 or Erk2, which share complete substrate redundancy (von Kriegsheim et al., 2009), abolished hypermotility with maintained base-line motility of DCs, indicating a dependency on both isoforms for hypermotility but not for base-line DC motility. This tight regulation is further supported by the abrogation of hypermotility upon knock down of Erk2 or Erk1, despite a compensatory elevation of Erk1 or Erk2 total protein expression, respectively. Moreover, the crucial dependence of DC hypermotility on Erk signaling cannot be generalized to other MAPK pathways, e.g. p38 MAPK signaling non-significantly impacted hypermotility of DCs (ten Hoeve et al., 2019). Together, these observations prompted us to investigate signaling pathways upstream of Erk activation in parasitized DCs.

Our data show that VGCC-CaM-CaMkII-Ras signaling mediate migratory activation of DCs upstream of Erk. Previously, we showed that the onset of the hypermigratory phenotype in DCs depends on live intracellular parasites and the discharge of *T. gondii* secretory organelles, independently of TLR-MyD88 signaling or chemotaxis (Fuks et al., 2012; Lambert et al., 2006; Olafsson et al., 2018; Sangare et al., 2019; Weidner et al., 2013). However, TLR-MyD88 signaling and chemotactic stimuli can also activate Erk. Specifically, in *T. gondii*-infected macrophages, both TLR-MyD88-dependent and a strong component of TLR-MyD88-independent Erk activation has been described (Kim et al., 2006). However, *T. gondii*-induced hypermigration is fully maintained in MyD88-deficient DCs (Lambert et al., 2006; Olafsson et al., 2018). Similarly, hypermigration is independent of chemotactic stimuli (but can act synergistically with chemotaxis upon stimulation) (Fuks et al., 2012; Kanatani et al., 2015; Weidner et al., 2013). Thus, alternative signaling underlies the migratory activation of DCs by *T. gondii* and, consequently, the hypermotility-related activation of Erk observed here.

*T. gondii* promotes DC hypermigration via GABAergic signaling (Fuks et al., 2012), which triggers the activation of the VGCC Ca_V_1.3 (Kanatani et al., 2017). VGCC-mediated Ca^2+^ influx impacts Ca^2+^-sensing proteins locally at the cell membrane (Rosen et al., 1994; Zhou et al., 2015). Consistent with this notion, our data show a crucial dependence of hypermigration on VGCC-CaM-CaMkII-Ras signaling. Here, by pharmacological agonism and antagonism of the VGCC signaling axis, we extend these findings by linking Erk phosphorylation to VGCC activation and CaM-CaMkII-Ras signaling. We show that inhibition of L-type VGCCs and Ca_V_1.3 with nifedipine and CPCPT, respectively, effectively blocks Erk phosphorylation in *T. gondii*-infected DCs. We previously showed that VGCC agonism enhanced migration of unchallenged and *T. gondii*-challenged DCs (Kanatani et al., 2017). Importantly, reinforcing the link between VGCCs/Ca_V_1.3 and Erk, we show that L-type VGCC agonism (Bay K) enhances Erk phosphorylation via CaM-CaMkII-Ras signaling while the opposite effect was observed by antagonism (nifedipine, CPCPT). These data are in line with paradigms in neuronal cells, where VGCC-dependent CaM activity regulates cytoskeleton organization and cell migration via Ras-Erk signaling (Rosen et al., 1994). Further, in lymphocytes, treatment with nifedipine and Bay K blocked and stimulated Erk phosphorylation, respectively (Kotturi et al., 2003). Thus, our data describe for the first time VGCC/Ca_V_1.3-mediated Erk activation in leukocytes and provide evidence for its dependence on CaM-CaMkII-Ras signaling.

Cancer cells seldom rely on one single pathway for orchestrating metastasis. Instead, metastasis generally relies on the activation of redundant and complementary pathways (Tracey et al., 2013). Consequently, we reasoned that diverse signaling cascades might underlie hypermotility in parasitized DCs.

Here, we identify a role for the receptor tyrosine kinase Met in the migratory activation of primary DCs. Upon *T. gondii* challenge, transcriptional upregulation and increased Met protein expression was indicative of activation. Importantly, antagonism of Met and gene silencing of *met* inhibited DC hypermotility. Functionally reinforcing the observation that Met is upregulated on the plasma membrane of parasitized DCs, recombinant Hgf (rHgf) treatment synergized with infection on migratory activation. Further, treatment with rHgf induced Erk phosphorylation and motility in unchallenged DCs. Conversely, Met antagonism inhibited Erk phosphorylation in unchallenged rHgf-treated DCs and *T. gondii*-infected DCs. Finally, Ras inhibition blocked rHgf-Met-induced Erk phosphorylation. Consistent with this idea, aberrant Met signaling mediates invasive cell growth and metastasis (Benvenuti and Comoglio, 2007) and afferent lymphatic migration of DC/Langerhans cells upon skin inflammation (Baek et al., 2012). Altogether, our data demonstrate that Met mediates migratory activation of DCs through Ras-Erk signaling.

However, the secreted amount of Met’s only known receptor ligand, Hgf, was non-significantly elevated in infected DC supernatants, opening up for the possibility of alternate mechanisms for Met activation. Ligand-independent activation of Met, or transactivation, is regulated by adapter proteins, integrins and tyrosine kinase cross-talk (Fischer et al., 2004; Hui et al., 2009; Mitra et al., 2011). In support for tyrosine kinase-mediated transactivation of Met, pharmacological antagonism of Fak and Pyk2 blocked Erk phosphorylation similar to Met antagonism in *T. gondii*-infected DCs. In breast cancer cells, the integrin Itgb1 transactivates Met (Barrow-McGee et al., 2016). We recently described a Timp-1-CD63-Itgb1-Fak signaling axis in *T. gondii*-infected DCs and gene silencing and antagonism of Itgb1 and Fak abolished hypermotility (Olafsson et al., 2019). Further, Timp-1 was recently shown to promote the activation of the RTK c-Kit in colorectal cancer (Nordgaard et al., 2019). Altogether, the data advocate for a potentiation of the motogenic action of Met through the Timp-1-CD63-Itgb1-Fak axis. However, additional, not mutually exclusive, possibilities exist. The secreted *T. gondii* proteins MIC1, 3 and 6 can directly phosphorylate the RTK Egfr (Muniz-Feliciano et al., 2013), which could hypothetically contribute to Met transactivation. Collectively, the data demonstrate that *T. gondii* activates the Met signaling pathway in infected DCs. Consistent with our previous studies, a milieu of RTK cross-talk with Erk phosphorylation in *T. gondii*-infected immune cells emerges, reminiscent of that described in metastatic cancer cells (Kaufmann et al., 2009; Volinsky and Kholodenko, 2013).

Based on the data at hand, we propose a model for the migratory activation of DCs upon challenge with *T. gondii* (Fig. 5): *T. gondii* infection activates Ras signaling through Met and GABA-VGCC-CaM-CaMkII signaling. Subsequently, the Ras-Raf-Mek cascade phosphorylates Erk, which drives migratory activation in DCs. The data show that *T. gondii* orchestrates the migratory activation of parasitized DCs through the Ras-Erk signaling node, which in turn is activated via tyrosine kinase signaling, integrins and GABA/Ca^2+^ ion channel activation (Kanatani et al., 2017; Olafsson et al., 2019). Our data also provide a molecular basis for the mesenchymal-to-amoeboid transition (MAT) and hypermigration of parasitized DCs. Interestingly, amoeboid motility is a feature of metastasizing cells and the signaling in parasitized DCs is consonant with emerging paradigms of signaling in metastatic cells (Lambert et al., 2017). Future studies will unravel the downstream effectors of Erk1/2 in DC hypermotility. In this context, putative parasite-derived effectors have been recently identified, which may interact with host cell MAPK signaling (Weidner et al., 2016), WAVE complex and actin dynamics (Sangare et al., 2019) and Rho GTPases (Drewry et al., 2019) in infected leukocytes. Thus, our data provide a signaling framework for assessing parasite-derived effectors and non-canonical migratory activation of leukocytes. Finally, this work highlights conserved signaling mechanisms between neurons, metastasizing cells and immune cells in the context of infection, which should be taken into consideration when designing therapies that target Met, VGCCs or Ras-Erk signaling.

**Figure 5.**
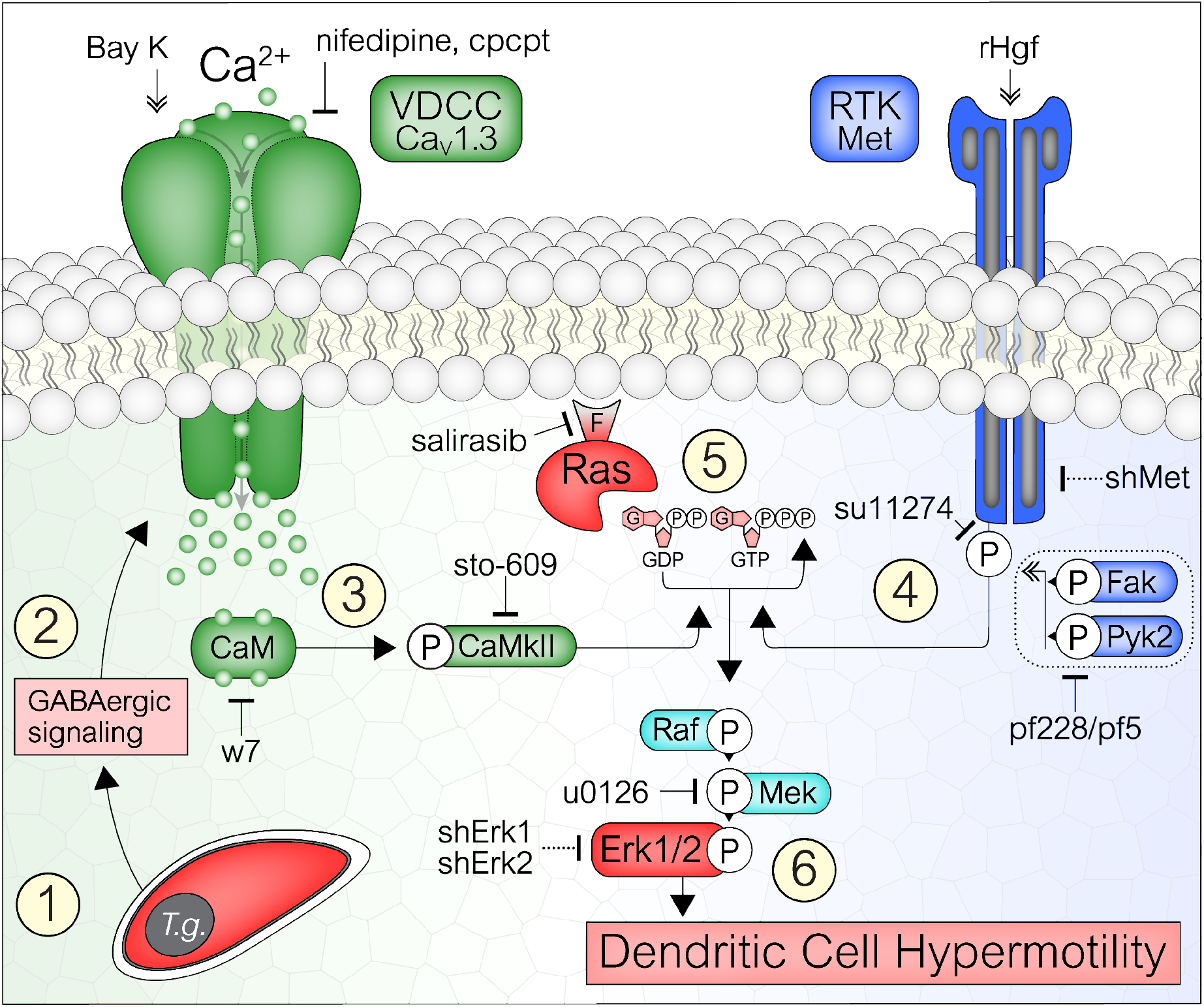
Model for Erk-mediated migratory activation of *T. gondii-infected* DCs via VGCC-Ras and Met-Ras signaling. **1.** *T. gondii* (*T.g.*) actively invades DCs and resides in a parasitophorous vacuole. **2.** Within 5 minutes of parasite invasion (Weidner et al., 2013), hypermotility is triggered in DCs through GABAergic signaling (Fuks et al., 2012). **3.** GABAergic signaling triggers membrane depolarization which activates VGCCs (chiefly Ca_V_1.3) and leads to Ca^2+^ influx. Gene silencing of Ca_V_1.3 (but not Ca_V_1.2) abrogates hypermotility (Kanatani et al., 2017). Inhibition of L-type VGCCs or Ca_V_1.3 with nifedipine and CPCPT, respectively, abolishes *T. gondii*-induced Erk phosphorylation and hypermotility. Similarly, inhibition of the VGCC signal transduction proteins CaM (w7) and CaMkII (sto-609) inhibit Erk phosphorylation and hypermotility. ‘P’ indicates protein phosphorylation. **4.** RTK Met expression in DCs is upregulated upon *T. gondii* infection and secretion of its ligand Hgf is maintained. Inhibition of Met phosphorylation (su11274) blocks Erk phosphorylation in *T. gondii*-infected DCs. Activation of Met with recombinant Hgf (rHgf) induces hypermotility and Erk phosphorylation in unchallenged and *T. gondii*-challenged DCs. Inhibition of Met phosphorylation (su11274) and gene silencing of met (shMet) abolish hypermotility. The integrin Itgb1 activates Fak upon *T. gondii* infection of DCs and gene silencing of Itgb1 and Fak inhibit hypermotility (Olafsson et al., 2019). Inhibition of the tyrosine kinases Fak and Pyk2 (pf228, pf5) blocks Erk phosphorylation, supporting tyrosine kinase-mediated transactivation of Met. **5.** VGCC-mediated Ca^2+^ influx and Met signaling converge on Ras GTPase. Antagonism of Ras farnesylation (indicated by ‘F’) with salirasib, inhibits Ras localization to the plasma membrane and its transition to activated form (GDP to GTP). Ras antagonism blocks agonist-induced (Bay K) VGCC-mediated Erk phosphorylation and hypermotility. Similarly, Ras antagonism blocks rHgf-induced Met-mediated Erk phosphorylation and hypermotility. **6.** Erk1/2 is rapidly phosphorylated in *T. gondii*-infected DCs downstream of Ras via Raf and Mek MAPKs. Antagonism of Mek (u0126) inhibits Erk phosphorylation and hypermotility in parasitized DCs. Further, gene silencing of Erk1 (shErk1) and Erk2 (shErk2) abolishes hypermotility. Hypothetically, activated Erk regulates targets in the cytosol and also translocates to the nucleus to regulate gene expression, to maintain the DC in a hypermotile state.

## Materials and Methods

### Ethics statement

The Regional Animal Research Ethical Board, Stockholm, Sweden, approved experimental procedures and protocols involving extraction of cells from mice, following proceedings described in EU legislation (Council Directive 2010/63/EU).

### Cells and Parasites

Murine bone marrow-derived DCs (DCs) were generated as previously described (Lambert et al., 2006). Briefly, cells from bone marrow of 6-10 week old male or female C57BL/6 mice (Charles River) were cultivated in RPMI 1640 with 10% fetal bovine serum (FBS), gentamicin (20 μg/ml), glutamine (2 mM) and HEPES (0.01 M), referred to as complete medium (CM; all reagents from Life Technologies), and supplemented with recombinant mouse GM-CSF (10 ng/ml) (Peprotech). Loosely adherent cells were harvested after 6 or 10 days of maturation. *T. gondii* tachyzoites of the RFP-expressing Prugniaud strain (Pru and Pru-RFP, type II) (Lambert et al., 2009) or GFP-expressing Ptg strain (type II) (Weidner et al., 2013) lines were maintained by serial 2-days passages in human foreskin fibroblast (HFF-1 SCRC-1041, American Type Culture Collection) monolayers. The hypermigratory phenotype induced by type II strains in DCs has been previously characterized (Olafsson et al., 2018).

### Reagents

The following soluble reagents were used to treat DCs in motility assays and for samples analyzed by western blotting: w-7 hydrochloride (Tocris), sto-609 acetate (Tocris), Hgf (40 ng/ml, Sigma), su11274 (Tocris), u0126 (Tocris), salirasib (Tocris), nifedipine (Sigma), Bay K8644 (Sigma).

### Lentiviral vector production and in vitro transduction

Self-complementary hairpin DNA oligos targeting mouse *met* (shMet), *erk1* (shErk1) *erk2* (shErk2) mRNA, and a non-related sequence shRNA control (shLuc) were chemically synthesized (DNA Technology, Denmark), aligned and ligated in a self-inactivating lentiviral vector (pLL3.7) containing a CMV-driven eGFP reporter and a U6 promoter upstream of cloning restriction sites (HpaI and XhoI) (Table S1). Lentivirus production was done using lipofectamine transfection. Briefly, shLuc, shMet, shErk1 or shErk2 vectors were co-transfected with pΔ8.91 packaging vector and pCMV-VSVg envelope vector into HEK293FT or HEK293T Lenti-X cells and the resulting supernatant was collected after 48-72 h. The supernatant was centrifuged to eliminate cell debris, filtered through 0.45-mm cellulose acetate filters, and virus particles were concentrated by ultracentrifugation and resuspended in RPMI. Titers were determined by infecting HEK293FT cells with serial dilutions of concentrated lentivirus and determination of percent of infected cells by flow cytometry (FACS Fortessa, BD). BMDCs were transduced twice (MOI 1, days 3 and 4) or once by spinoculation at 1200xg for 2 h (MOI 2, day 3). Three days post transduction eGFP-expression was verified by epifluorescence microscopy before the cells were used in experiments.

### Motility assays

Motility assays were performed as previously described (Olafsson et al., 2019). Briefly, DCs were cultured in 96-well plates with CM ± freshly egressed *T. gondii* tachyzoites (Pru-RFP for shRNA experiments and Ptg-GFP for all other experiments, MOI 3, 4 h) and soluble reagents as indicated. Bovine collagen I (1mg/ml, Life Technologies) was then added and live cell imaging was performed for 1 h, 1 frame/ min, at 10x magnification (Z1 Observer with Zen 2 Blue v. 4.0.3, Zeiss). Time-lapse images were consolidated into stacks and motility data was obtained from >30 cells/condition (Manual Tracking, ImageJ) yielding mean velocities (Chemotaxis and migration tool, v. 2.0). Infected cells were defined by RFP- or GFP-cell co-localization. Transduced cells were defined by eGFP or tGFP reporter expression.

### Quantitively polymerase chain reaction (qPCR)

DCs were cultured in CM ± freshly egressed *T. gondii* tachyzoites (Ptg-GFP, MOI 3), at indicated time points. Total RNA was extracted using TRIzol reagent (Invitrogen). qPCR was performed using SYBR Green PCR master mix (Kapa biosystems), forward and reverse primers (200 nM) and cDNA (100 ng) and run on the Rotor Gene 6000 system (Corbett) for 45 cycles. Amplicons were validated with melting curves and agarose gel (1%) analysis and data was analyzed with RG-6000 application software (v1.7, Corbett). *Gapdh* and *actb* were used as housekeeping genes to generate ΔCt values. 2^(-ΔCt)^ values were generated to show relative knockdown efficiency and 2^(-ΔΔCt)^ values were used to show expression fold-change upon infection. Fold changes in expression were calculated using the comparative DCT method against the non-infected DC control. Primers were designed using Getprime (Table S2) and purchased from Invitrogen.

### Flow cytometry

DCs were cultured in CM ± freshly egressed *T. gondii* tachyzoites (Ptg-GFP, MOI 3, 4 h). Following Fc blockade (clone 2.4G2, BD Pharmingen), cells were stained with anti-CD11c-PE-Cy7 (clone N418, eBiosciences). After fixation (PFA 4%) cells were either directly stained (plasma membrane Met) with anti-Met-PE (clone eBioclone 7, eBiosciences) or isotype control (clone eBRG1, eBiosciences) or permeabilized (IntraPrep Permeabilization kit, Beckman Coulter) before staining (total Met). All antibodies were used at a dilution of 1:100. Samples were run on the BD LSRFortessa Cell Analyzer (BD). Data was analyzed in FlowJo (Tree Star Inc, OR). For infection frequency, DCs were cultured in CM ± reagents (as indicated) and freshly egressed *T. gondii* tachyzoites (Ptg-GFP, MOI 3) for 2 h. After fixation (PFA 4%), samples were run with the easyCyte^™^ 8HT (Millipore), in accordance with the manufacturer’s guidelines. Samples were gated on FSC, SSC and GFP. A minimum of 30.000 cells were analyzed per condition. Data were analyzed in FlowJo (Tree Star Inc, OR).

### Hgf ELISA

Supernatants from DCs cultured in CM ± freshly egressed *T. gondii* tachyzoites (Ptg-GFP, MOI 3) were collected 2, 6, 12 or 24 h post infection. Samples were analyzed by ELISA (MHG00, ThermoFisher) according to manufacturer’s instruction. Samples were normalized to a CM control.

### Western Blotting

Cell lysates were collected and sonicated in RIPA buffer (150mM NaCl, 5mM EDTA, pH 8.0, 50 mM Tris, pH 8.0, 0.1% Triton-X100, 0.5% sodium deoxycholate, 0.1% SDS) with protease and phosphatase inhibitors (Thermo Scientific). Proteins were separated on 10% SDS-PAGE gels followed by transfer to polyvinylidene difluoride Immobilon membranes (Millipore). Phospho-Erk was detected using a phospho-Erk antibody (clone 4695P, Cell Signaling), total Erk was detected using an Erk antibody (clone 4370S, Cell Signaling) and Gapdh was detected with (clone ABS16, Millipore). Anti-Rabbit-HRP (clone 7074S, Cell Signaling) was used as secondary antibody. Proteins were visualized using ECL reagents (GE healthcare) in a Biorad ChemiDoc XRS^+^. Densitometry analysis was performed using image J (NIH, MD, USA).

### Statistical analyses

Multiple comparisons of normally distributed data were carried out with one-way ANOVA followed by Tukey HSD or Holm-Sidak’s multiple comparisons test. Two-way ANOVA followed by Holm-Sidak’s multiple comparisons test was used to evaluate the interaction between time and infection. Two sample comparisons were carried out with Paired *t-*test. Normality was tested by the Shapiro-Wilks test. In all statistical tests p-values ≥ 0.05 were defined as non-significant and p-values < 0.05 were defined as significant. All statistics were performed with Prism (v. 8, Graphpad).

## Acknowledgements

This work was funded by the Swedish Research Council (Vetenskapsrådet, 2018-02411 to A.B.) and the Olle Engkvist Foundation (193-609 to A.B.) We thank all members of the Barragan lab for critical input.

## Supporting information

**S1 Table.**
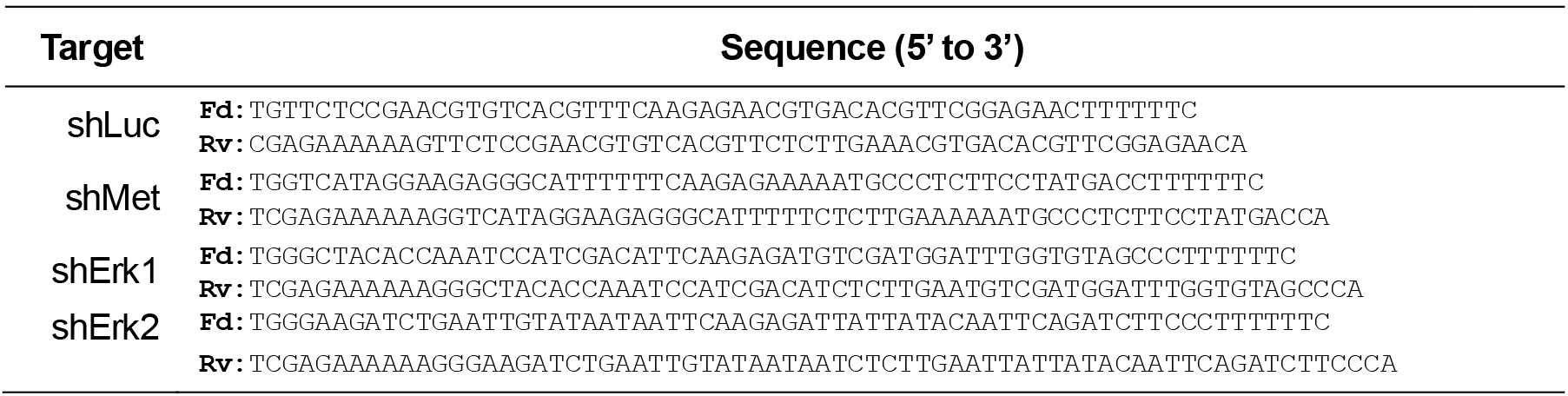
Sequences of shRNAs.

**S2 Table.**
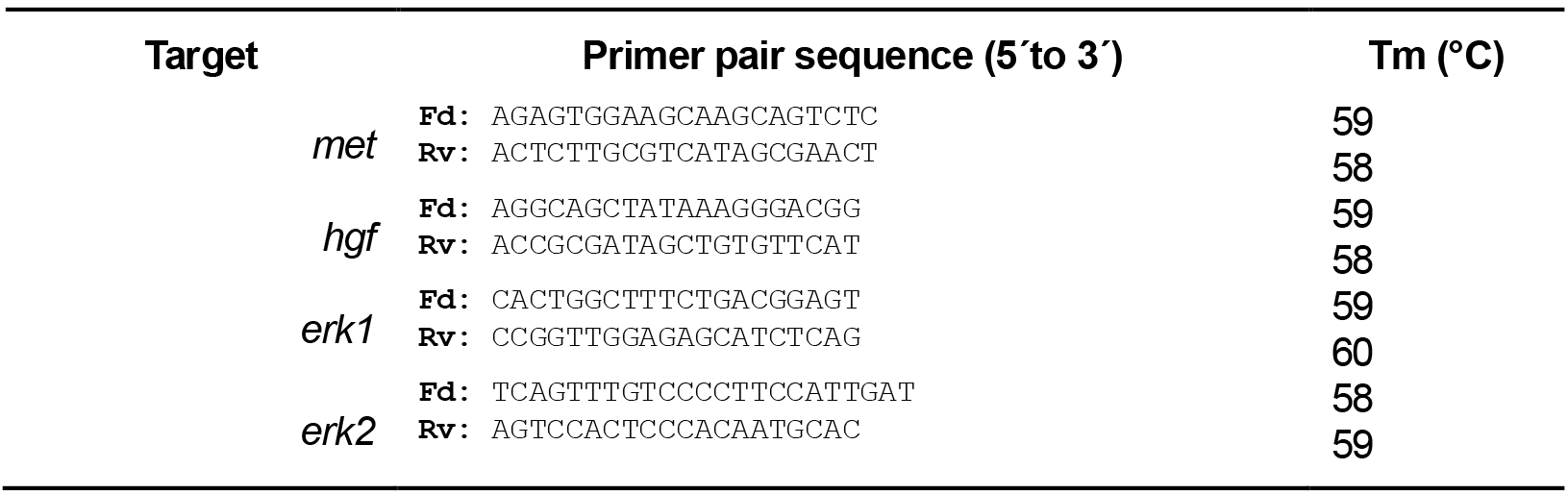
PCR primers used to amplify *met, hgf, erk1* and *erk2*.

**Figure S1.**
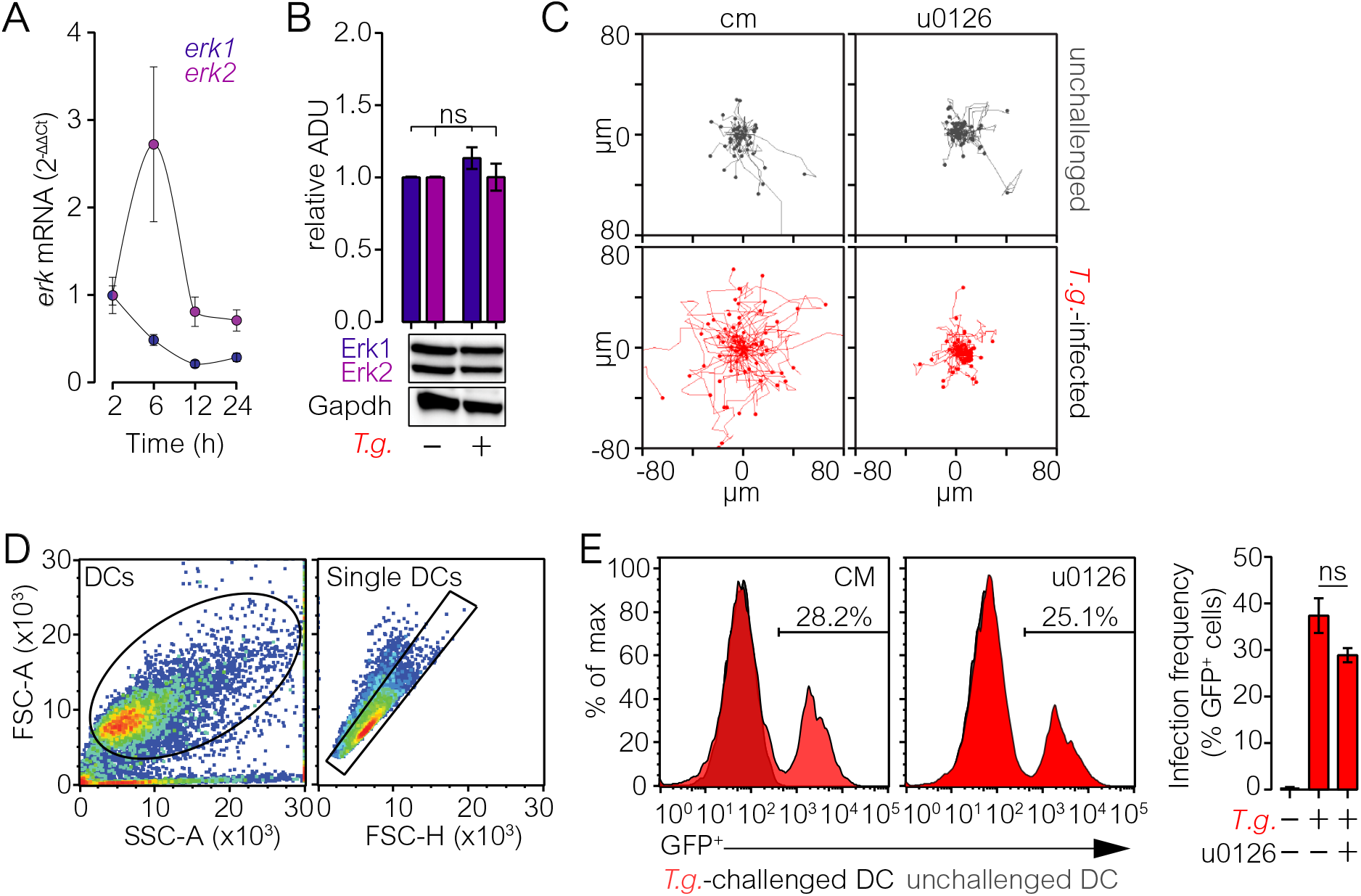
Expression of Erk1/2 by *T. gondii*-challenged DCs, phosphorylation of Erk in unchallenged DCs, impact of Erk inhibition on DC motility and infection frequencies. **A.** Fold change in *erk1* and *erk2* mRNA expression in *T. gondii*-challenged DCs at 2, 6, 12 and 24 h post infection (hpi). Graph shows mean expression (± SEM) related to gapdh (2^-ΔCt^) and unchallenged DCs (2^-ΔΔCt^, n=3 independent experiments). **B.** Total Erk protein expression in unchallenged and *T. gondii*-challenged DCs at 6 hpi. Bar graph shows mean (± SEM) Erk total protein related to Gapdh. Expression of unchallenged DCs was set to 1. Images show representative blots used for quantification from 1 experiment (n=4 biological replicates from independent blots). **C.** Representative motility plots of unchallenged and *T. gondii*-infected DCs embedded in collagen ± u0126 (10 μM). Plots are representative of 3 independent experiments. **D.** Representative dot plots of flow cytometry analyses of *T. gondii* infection frequency. Dot plots show gating of cells (FSC-A/ SSC-A) and single cells (FSC-A/FSC-H). **E.** *T. gondii* infection frequencies of DCs cultured in complete medium (cm) ± u0126 (10 μM). Histograms show gates used to define *T. gondii* (GFP+)-infected cells in unchallenged (grey fill) and *T. gondii*-challenged (red fill) DCs and are representative of 3 experiments. Bar graph shows mean percentage (± SEM) of GFP+ cells (n=3). Ns indicates non-significant difference: One-way ANOVA, Tukey’s HSD post-hoc test (B and E).

**Figure S2.**
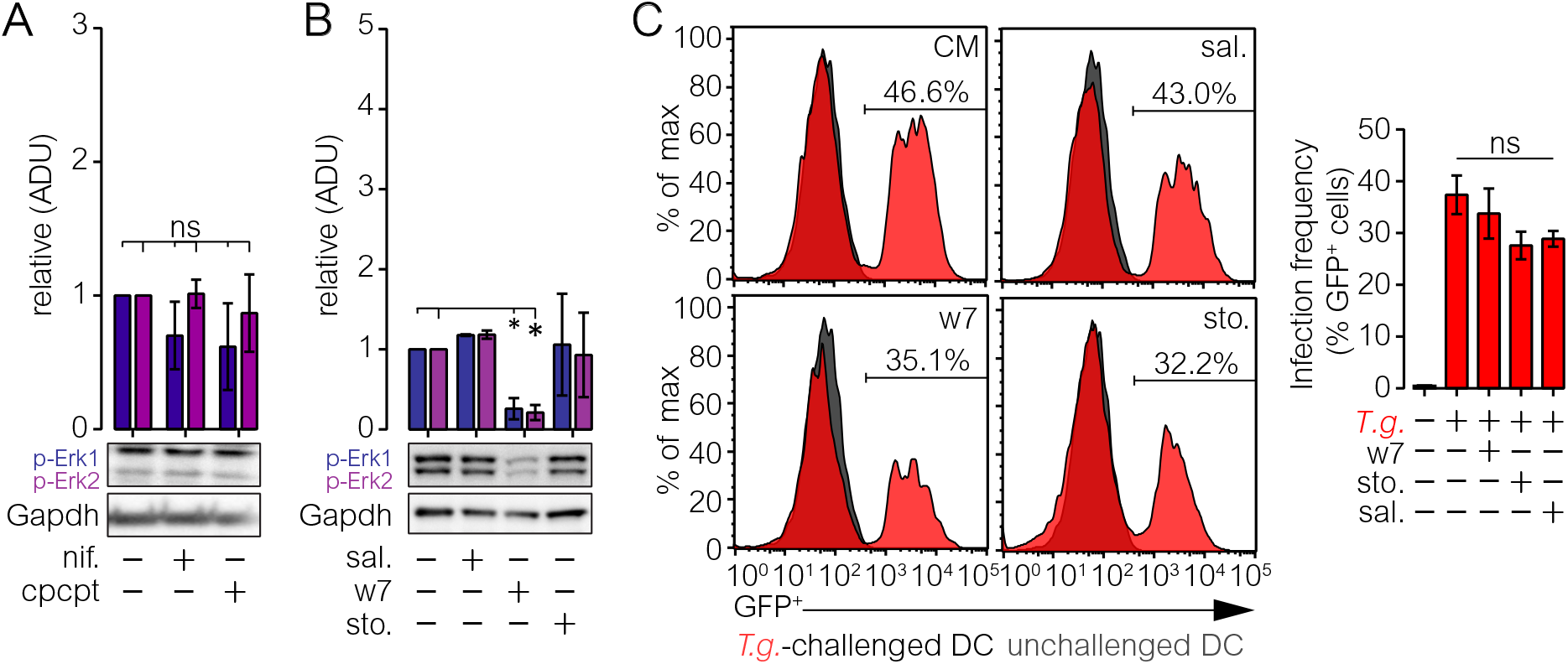
Erk phosphorylation in unchallenged DCs in presence of VGCC-CaM-CaMkII-Ras inhibitors and infection frequencies. **A.** Erk phosphorylation in unchallenged DCs ± nifedipine (50 μM) or CPCPT (10 μM) treatment 2 hpi. Bar graph shows mean (± SEM) phosphorylated Erk1 and Erk2 protein related to Gapdh. Expression of unchallenged DCs was set to 1. Images show representative blots used for quantification from 1 experiment (n=3 biological replicates from independent blots). **B.** Erk phosphorylation in unchallenged DCs ± salirasib (100 μM), w7 (25 μM) or sto-609 (sto., 50 μM) treatment 2 hpi. Bar graph shows mean (± SEM) phosphorylated Erk1 and Erk2 protein related to Gapdh. Expression of unchallenged DCs was set to 1. Images show representative blots used for quantification from 1 experiment (n=3 biological replicates from independent blots). **C.** *T. gondii* infection frequency of DCs cultured in cm ± salirasib (sal.,100 μM), w7 (25 μM) or sto-609 (sto., 50 μM). Histograms show gates used to define *T. gondii* (GFP+)-infected cells in unchallenged (grey fill) and *T. gondii*-challenged (red fill) DCs and are representative of 3 experiments. Bar graph shows mean percentage (± SEM) of GFP+ cells (n=3). Asterisks (*) indicate significant difference, ns: non-significant difference: One-way ANOVA, Tukey’s HSD post-hoc test (A and C), Unpaired t-test (B).

**Figure S3.**
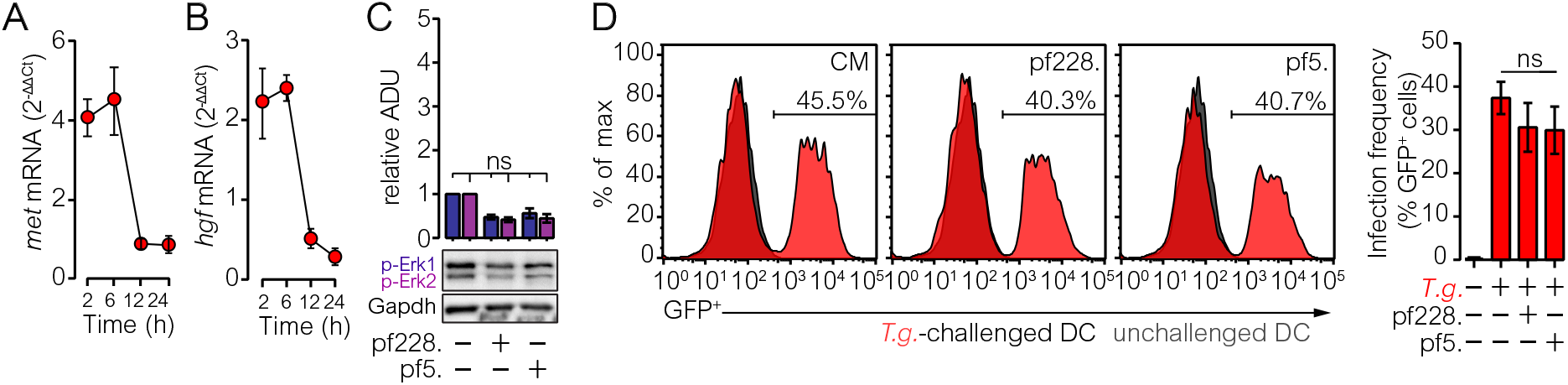
Met and hgf expression in *T. gondii*-challenged DCs, impact of Fak inhibition on Erk phosphorylation in unchallenged DCs and infection frequencies. **A.** Fold change Met mRNA expression in *T. gondii*-challenged DCs at 2, 6, 12 and 24 hpi. Bar graph shows mean expression (± SEM) related to gapdh (2^-ΔCt^) and unchallenged DCs (2^-ΔΔCt^, n=3 independent experiments). **B.** Fold change Hgf mRNA expression in *T. gondii*-challenged DCs at 2, 6, 12 and 24 hpi. Bar graph shows mean expression (± SEM) related to gapdh (2^-ΔCt^) and unchallenged DCs (2^-ΔΔCt^, n=3 independent experiments). **C.** Erk phosphorylation in unchallenged DCs ± pf228 (10 μM) or pf5 (10 μM) treatment 2 hpi. Bar graph shows mean (± SEM) phosphorylated Erk1 and Erk2 protein related to Gapdh. Expression of unchallenged DCs was set to 1. Images show representative blots used for quantification from 1 experiment (n=3 biological replicates from independent blots). **D.** *T. gondii* infection frequency of DCs cultured in cm ± pf228 (10 μM) or pf5 (10 μM). Histograms show gates used to define *T. gondii* (GFP+)-infected cells in unchallenged (grey fill) and *T. gondii*-challenged (red fill) DCs and are representative of 3 experiments. Bar graph shows mean percentage (± SEM) of GFP^+^ cells (n=3). ns: non-significant difference: One-way ANOVA, Holm-Sidak’s multiple comparisons test (C) or Tukey’s HSD post-hoc test (D).

**Figure S4.**
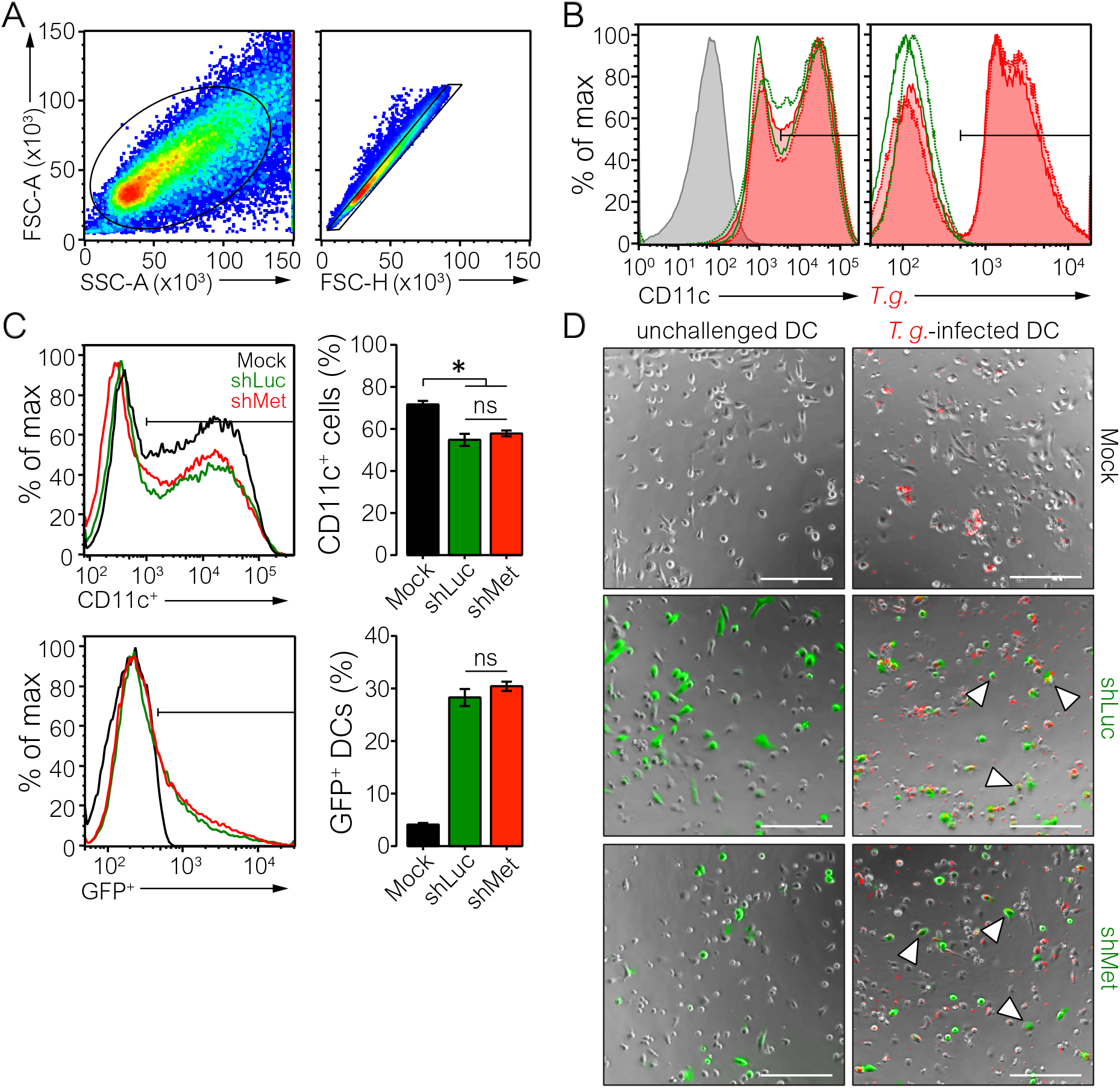
Characterizations of Met expression and gene silencing of met for motility analyses. **A.** Representative dot plots used in flow cytometry analyses of Met protein expression. Dot plots show gating of cells (FSC-A/SSC-A) and single cells (FSC-A/ FSC-H). Plots are representative of 3 experiments. **B.** Histograms show gates used to define CD11c^+^ cells (left) and GFP^+^ cells (*T.g.*; right) in unchallenged (green) and *T. gondii*-infected (red) cells used in Met protein analyses. Dashed and continuous lines show signal corresponding to total Met (permeabilized) and plasma membrane Met (unpermeabilized), respectively. Grey fill represents unstained control for CD11c. Histograms are representative of 3 experiments. **C.** Histograms show gates used to define CD11c^+^ cells (top) and GFP^+^cells (bottom) in mock-treated DCs (Mock, black), shMet (red) and shLuc (green) transduced cells. Bar graph shows mean (± SEM) percentage of CD11c^+^ (top) and GFP+ (bottom) cells for each condition (n=3). **D.** Representative micrographs of mock-treated DCs (Mock) and eGFP-expressing DCs transduced with lentiviral vectors targeting met mRNA (shMet) or a non-expressed target (shLuc) embedded in collagen ± RFP-expressing *T. gondii* tachyzoites. Arrowheads indicate *T. gondii*-infected cells expressing an eGFP-reporter assessed in the assay. Scale bars: 200 μm. Micrographs are representative of 3 independent experiments. Asterisk (*) indicates significant difference, ns: non-significant difference: One-way ANOVA, Tukey’s HSD post-hoc test (C)

## References

Agell, N., O. Bachs, N. Rocamora, and P. Villalonga. 2002. Modulation of the Ras/Raf/MEK/ERK pathway by Ca(2+), and calmodulin. Cell Signal. 14:649–654.

Baek, J.H., C. Birchmeier, M. Zenke, and T. Hieronymus. 2012. The HGF receptor/Met tyrosine kinase is a key regulator of dendritic cell migration in skin immunity. J Immunol. 189:1699–1707.

Barrow-McGee, R., N. Kishi, C. Joffre, L. Menard, A. Hervieu, B.A. Bakhouche, A.J. Noval, A. Mai, C. Guzman, L. Robbez-Masson, X. Iturrioz, J. Hulit, C.H. Brennan, I.R. Hart, P.J. Parker, J. Ivaska, and S. Kermorgant. 2016. Beta 1-integrin-c-Met cooperation reveals an inside-in survival signalling on autophagy-related endomembranes. Nat Commun. 7:11942.

Benvenuti, S., and P.M. Comoglio. 2007. The MET receptor tyrosine kinase in invasion and metastasis. Journal of cellular physiology. 213:316–325.

Charest, P.G., Z. Shen, A. Lakoduk, A.T. Sasaki, S.P. Briggs, and R.A. Firtel. 2010. A Ras signaling complex controls the RasC-TORC2 pathway and directed cell migration. Dev Cell. 18:737–749.

Chernyavsky, A.I., J. Arredondo, E. Karlsson, I. Wessler, and S.A. Grando. 2005. The Ras/Raf-1/MEK1/ERK signaling pathway coupled to integrin expression mediates cholinergic regulation of keratinocyte directional migration. J Biol Chem. 280:39220–39228.

Courret, N., S. Darche, P. Sonigo, G. Milon, D. Buzoni-Gatel, and I. Tardieux. 2006. CD11c- and CD11b-expressing mouse leukocytes transport single Toxoplasma gondii tachyzoites to the brain. Blood. 107:309–316.

Cox, A.D., M.M. Hisaka, J.E. Buss, and C.J. Der. 1992. Specific isoprenoid modification is required for function of normal, but not oncogenic, Ras protein. Mol Cell Biol. 12:2606–2615.

Dolmetsch, R.E., U. Pajvani, K. Fife, J.M. Spotts, and M.E. Greenberg. 2001. Signaling to the nucleus by an L-type calcium channel-calmodulin complex through the MAP kinase pathway. Science. 294:333–339.

Drewry, L.L., N.G. Jones, Q. Wang, M.D. Onken, M.J. Miller, and L.D. Sibley. 2019. The secreted kinase ROP17 promotes Toxoplasma gondii dissemination by hijacking monocyte tissue migration. Nat Microbiol. 4:1951–1963.

Ebert, P.J.R., J. Cheung, Y. Yang, E. McNamara, R. Hong, M. Moskalenko, S.E. Gould, H. Maecker, B.A. Irving, J.M. Kim, M. Belvin, and I. Mellman. 2016. MAP Kinase Inhibition Promotes T Cell and Anti-tumor Activity in Combination with PD-L1 Checkpoint Blockade. Immunity. 44:609–621.

Eblen, S.T. 2018. Extracellular-Regulated Kinases: Signaling From Ras to ERK Substrates to Control Biological Outcomes. Adv Cancer Res. 138:99–142.

Fischer, O.M., S. Giordano, P.M. Comoglio, and A. Ullrich. 2004. Reactive oxygen species mediate Met receptor transactivation by G protein-coupled receptors and the epidermal growth factor receptor in human carcinoma cells. J Biol Chem. 279:28970–28978.

Fuks, J.M., R.B. Arrighi, J.M. Weidner, S. Kumar Mendu, Z. Jin, R.P. Wallin, B. Rethi, B. Birnir, and A. Barragan. 2012. GABAergic signaling is linked to a hypermigratory phenotype in dendritic cells infected by Toxoplasma gondii. PLoS Pathog. 8:e1003051.

Giehl, K., B. Skripczynski, A. Mansard, A. Menke, and P. Gierschik. 2000. Growth factor-dependent activation of the Ras-Raf-MEK-MAPK pathway in the human pancreatic carcinoma cell line PANC-1 carrying activated K-ras: implications for cell proliferation and cell migration. Oncogene. 19:2930–2942.

Hartmann, G., K.M. Weidner, H. Schwarz, and W. Birchmeier. 1994. The motility signal of scatter factor/hepatocyte growth factor mediated through the receptor tyrosine kinase met requires intracellular action of Ras. J Biol Chem. 269:21936–21939.

Hui, A.Y., J.A. Meens, C. Schick, S.L. Organ, H. Qiao, E.A. Tremblay, E. Schaeffer, S. Uniyal, B.M. Chan, and B.E. Elliott. 2009. Src and FAK mediate cell-matrix adhesion-dependent activation of Met during transformation of breast epithelial cells. J Cell Biochem. 107:1168–1181.

Illario, M., A.L. Cavallo, K.U. Bayer, T. Di Matola, G. Fenzi, G. Rossi, and M. Vitale. 2003. Calcium/calmodulin-dependent protein kinase II binds to Raf-1 and modulates integrin-stimulated ERK activation. The Journal of biological chemistry. 278:45101–45108.

Kanatani, S., J.M. Fuks, E.B. Olafsson, L. Westermark, B. Chambers, M. Varas-Godoy, P. Uhlen, and A. Barragan. 2017. Voltage-dependent calcium channel signaling mediates GABAA receptor-induced migratory activation of dendritic cells infected by Toxoplasma gondii. PLoS Pathog. 13:e1006739.

Kanatani, S., P. Uhlen, and A. Barragan. 2015. Infection by Toxoplasma gondii Induces Amoeboid-Like Migration of Dendritic Cells in a Three-Dimensional Collagen Matrix. PloS one. 10:e0139104.

Kaufmann, R., C. Oettel, A. Horn, K.J. Halbhuber, A. Eitner, R. Krieg, K. Katenkamp, P. Henklein, M. Westermann, F.D. Bohmer, R. Ramachandran, M. Saifeddine, M.D. Hollenberg, and U. Settmacher. 2009. Met receptor tyrosine kinase transactivation is involved in proteinase-activated receptor-2-mediated hepatocellular carcinoma cell invasion. Carcinogenesis. 30:1487–1496.

Kim, L., B.A. Butcher, C.W. Lee, S. Uematsu, S. Akira, and E.Y. Denkers. 2006. Toxoplasma gondii genotype determines MyD88-dependent signaling in infected macrophages. J Immunol. 177:2584–2591.

Kotturi, M.F., D.A. Carlow, J.C. Lee, H.J. Ziltener, and W.A. Jefferies. 2003. Identification and functional characterization of voltage-dependent calcium channels in T lymphocytes. The Journal of biological chemistry. 278:46949–46960.

Lambert, A.W., D.R. Pattabiraman, and R.A. Weinberg. 2017. Emerging Biological Principles of Metastasis. Cell. 168:670–691.

Lambert, H., N. Hitziger, I. Dellacasa, M. Svensson, and A. Barragan. 2006. Induction of dendritic cell migration upon Toxoplasma gondii infection potentiates parasite dissemination. Cell Microbiol. 8:1611–1623.

Lambert, H., P.P. Vutova, W.C. Adams, K. Lore, and A. Barragan. 2009. The Toxoplasma gondii-shuttling function of dendritic cells is linked to the parasite genotype. Infect Immun. 77:1679–1688.

Lemmon, M.A., and J. Schlessinger. 2010. Cell signaling by receptor tyrosine kinases. Cell. 141:1117–1134.

Li, Y., J.L. Lin, R.S. Reiter, K. Daniels, D.R. Soll, and J.J. Lin. 2004. Caldesmon mutant defective in Ca(2+)-calmodulin binding interferes with assembly of stress fibers and affects cell morphology, growth and motility. J Cell Sci. 117:3593–3604.

Liu, C.H., Y.T. Fan, A. Dias, L. Esper, R.A. Corn, A. Bafica, F.S. Machado, and J. Aliberti. 2006. Cutting edge: dendritic cells are essential for in vivo IL-12 production and development of resistance against Toxoplasma gondii infection in mice. J Immunol. 177:31–35.

Lundberg, M.S., K.A. Curto, C. Bilato, R.E. Monticone, and M.T. Crow. 1998. Regulation of vascular smooth muscle migration by mitogen-activated protein kinase and calcium/calmodulin-dependent protein kinase II signaling pathways. J Mol Cell Cardiol. 30:2377–2389.

Mitra, A.K., K. Sawada, P. Tiwari, K. Mui, K. Gwin, and E. Lengyel. 2011. Ligand-independent activation of c-Met by fibronectin and alpha(5)beta(1)-integrin regulates ovarian cancer invasion and metastasis. Oncogene. 30:1566–1576.

Montoya, J.G., and O. Liesenfeld. 2004. Toxoplasmosis. Lancet. 363:1965–1976.

Muniz-Feliciano, L., J. Van Grol, J.A. Portillo, L. Liew, B. Liu, C.R. Carlin, V.B. Carruthers, S. Matthews, and C.S. Subauste. 2013. Toxoplasma gondii-induced activation of EGFR prevents autophagy protein-mediated killing of the parasite. PLoS Pathog. 9:e1003809.

Nordgaard, C., S. Doll, A. Matos, M. Hoeberg, J.U. Kazi, S. Friis, J. Stenvang, L. Ronnstrand, M. Mann, and J.M.A. Moreira. 2019. Metallopeptidase Inhibitor 1 (TIMP-1) promotes receptor tyrosine kinase c-Kit signaling in colorectal cancer. Mol Oncol.

Olafsson, E.B., E.C. Ross, M. Varas-Godoy, and A. Barragan. 2019. TIMP-1 promotes hypermigration of Toxoplasma-infected primary dendritic cells via CD63-ITGB1-FAK signaling. J Cell Sci. 132.

Olafsson, E.B., M. Varas-Godoy, and A. Barragan. 2018. Toxoplasma gondii infection shifts dendritic cells into an amoeboid rapid migration mode encompassing podosome dissolution, secretion of TIMP-1, and reduced proteolysis of extracellular matrix. Cell Microbiol. 20.

Pappas, G., N. Roussos, and M.E. Falagas. 2009. Toxoplasmosis snapshots: global status of Toxoplasma gondii seroprevalence and implications for pregnancy and congenital toxoplasmosis. International journal for parasitology. 39:1385–1394.

Riegel, K., J. Schloder, M. Sobczak, H. Jonuleit, B. Thiede, H. Schild, and K. Rajalingam. 2019. RAF kinases are stabilized and required for dendritic cell differentiation and function. Cell Death Differ.

Rosen, L.B., D.D. Ginty, M.J. Weber, and M.E. Greenberg. 1994. Membrane depolarization and calcium influx stimulate MEK and MAP kinase via activation of Ras. Neuron. 12:1207–1221.

Sangare, L.O., E.B. Olafsson, Y. Wang, N. Yang, L. Julien, A. Camejo, P. Pesavento, S.M. Sidik, S. Lourido, A. Barragan, and J.P.J. Saeij. 2019. In Vivo CRISPR Screen Identifies TgWIP as a Toxoplasma Modulator of Dendritic Cell Migration. Cell host & microbe. 26:478–492 e478.

Sasaki, A.T., C. Chun, K. Takeda, and R.A. Firtel. 2004. Localized Ras signaling at the leading edge regulates PI3K, cell polarity, and directional cell movement. J Cell Biol. 167:505–518.

Scheele, J.S., R.E. Marks, and G.R. Boss. 2007. Signaling by small GTPases in the immune system. Immunol Rev. 218:92–101.

Schlaepfer, D.D., S.K. Hanks, T. Hunter, and P. van der Geer. 1994. Integrin-mediated signal transduction linked to Ras pathway by GRB2 binding to focal adhesion kinase. Nature. 372:786–791.

Schwanhausser, B., D. Busse, N. Li, G. Dittmar, J. Schuchhardt, J. Wolf, W. Chen, and M. Selbach. 2011. Global quantification of mammalian gene expression control. Nature. 473:337–342.

Shi, S., J. Zhang, M. Liu, H. Dong, and N. Li. 2019. Ras-ERK signalling represses H1.4 phosphorylation at serine 36 to promote non-small-cell lung carcinoma cells growth and migration. Artif Cells Nanomed Biotechnol. 47:2343–2351.

Sibley, L.D. 2004. Intracellular parasite invasion strategies. Science. 304:248–253.

Stupack, D.G., S.Y. Cho, and R.L. Klemke. 2000. Molecular signaling mechanisms of cell migration and invasion. Immunol Res. 21:83–88.

ten Hoeve, A.L., M.-A. Hakimi, and A. Barragan. 2019. Sustained Egr-1 Response via p38 MAP Kinase Signaling Modulates Early Immune Responses of Dendritic Cells Parasitized by Toxoplasma gondii. Frontiers in Cellular and Infection Microbiology. 9.

Tracey, A., T.A. Martin, A.J. Sanders, J. Lane, and W.G. Jiang. 2013. Cancer Invasion and Metastasis: Molecular and Cellular Perspective. Landes Bioscience, Austin (TX).

Viala, E., and J. Pouyssegur. 2004. Regulation of tumor cell motility by ERK mitogen-activated protein kinases. Ann N Y Acad Sci. 1030:208–218.

Volinsky, N., and B.N. Kholodenko. 2013. Complexity of receptor tyrosine kinase signal processing. Cold Spring Harb Perspect Biol. 5:a009043.

von Kriegsheim, A., D. Baiocchi, M. Birtwistle, D. Sumpton, W. Bienvenut, N. Morrice, K. Yamada, A. Lamond, G. Kalna, R. Orton, D. Gilbert, and W. Kolch. 2009. Cell fate decisions are specified by the dynamic ERK interactome. Nat Cell Biol. 11:1458–1464.

Wang, D., Z. Li, E.M. Messing, and G. Wu. 2002. Activation of Ras/Erk pathway by a novel MET-interacting protein RanBPM. J Biol Chem. 277:36216–36222.

Webb, C.P., G.A. Taylor, M. Jeffers, M. Fiscella, M. Oskarsson, J.H. Resau, and G.F. Vande Woude. 1998. Evidence for a role of Met-HGF/SF during Ras-mediated tumorigenesis/metastasis. Oncogene. 17:2019–2025.

Weidner, J.M., and A. Barragan. 2014. Tightly regulated migratory subversion of immune cells promotes the dissemination of Toxoplasma gondii. Int J Parasitol. 44:85–90.

Weidner, J.M., S. Kanatani, M.A. Hernandez-Castaneda, J.M. Fuks, B. Rethi, R.P. Wallin, and A. Barragan. 2013. Rapid cytoskeleton remodelling in dendritic cells following invasion by Toxoplasma gondii coincides with the onset of a hypermigratory phenotype. Cell Microbiol. 15:1735–1752.

Weidner, J.M., S. Kanatani, H. Uchtenhagen, M. Varas-Godoy, T. Schulte, K. Engelberg, M.J. Gubbels, H.S. Sun, R.E. Harrison, A. Achour, and A. Barragan. 2016. Migratory activation of parasitized dendritic cells by the protozoan Toxoplasma gondii 14-3-3 protein. Cellular microbiology. 18:1537–1550.

Zhou, Y., C.O. Wong, K.J. Cho, D. van der Hoeven, H. Liang, D.P. Thakur, J. Luo, M. Babic, K.E. Zinsmaier, M.X. Zhu, H. Hu, K. Venkatachalam, and J.F. Hancock. 2015. SIGNAL TRANSDUCTION. Membrane potential modulates plasma membrane phospholipid dynamics and K-Ras signaling. Science. 349:873–876.

